# High-dimensional Ageome Representations of Biological Aging across Functional Modules

**DOI:** 10.1101/2024.09.17.613599

**Authors:** Kejun Ying, Alexander Tyshkovskiy, Qingwen Chen, Eric Latorre-Crespo, Bohan Zhang, Hanna Liu, Benyamin Matei-Dediu, Jesse R. Poganik, Mahdi Moqri, Kristina Kirschne, Jessica Lasky-Su, Vadim N. Gladyshev

## Abstract

The aging process involves numerous molecular changes that lead to functional decline and increased disease and mortality risk. While epigenetic aging clocks have shown accuracy in predicting biological age, they typically provide single estimates for the samples and lack mechanistic insights. In this study, we challenge the paradigm that aging can be sufficiently described with a single biological age estimate. We describe Ageome, a computational framework for measuring the epigenetic age of thousands of molecular pathways simultaneously in mice and humans. Ageome is based on the premise that an organism’s overall biological age can be approximated by the collective ages of its functional modules, which may age at different rates and have different biological ages. We show that, unlike conventional clocks, Ageome provides a high-dimensional representation of biological aging across cellular functions, enabling comprehensive assessment of aging dynamics within an individual, in a population, and across species. Application of Ageome to longevity intervention models revealed distinct patterns of pathway-specific age deceleration. Notably, cell reprogramming, while rejuvenating cells, also accelerated aging of some functional modules. When applied to human cohorts, Ageome demonstrated heterogeneity in predictive power for mortality risk, and some modules showed better performance in predicting the onset of age-related diseases, especially cancer, compared to existing clocks. Together, the Ageome framework offers a comprehensive and interpretable approach for assessing aging, providing insights into mechanisms and targets for intervention.

## Introduction

Aging is a complex biological process characterized by the progressive accumulation of molecular and cellular damage, leading to functional decline across various organ systems and ultimately increased mortality risk ^1^. While the precise mechanisms underlying aging are not fully elucidated, substantial evidence points to the critical role of epigenetic alterations in this process ^2^. Among these epigenetic modifications, DNA methylation has emerged as a key area of focus in aging research.

In mammalian systems, DNA methylation primarily occurs as 5-methylcytosine (5mC), a modification catalyzed by DNA methyltransferases (DNMTs) ^3,4^. Age-associated changes in DNA methylation patterns have been well-documented, with studies revealing a general trend of global hypomethylation accompanied by localized regions of hypermethylation ^5–8^. These age-related methylation changes exhibit remarkable consistency across individuals, enabling the development of highly accurate predictive models known as ‘epigenetic aging clocks’ ^9–11^. These clocks have demonstrated a stronger correlation with various health metrics compared to chronological age, suggesting their potential as more accurate indicators of biological age ^12,13^. However, despite their predictive power, these tools face limitations with regard to interpretability. Most notably, they typically provide a single estimate of biological age for the entire sample, tissue or organism, potentially overlooking heterogeneity in aging rates across functional modules within the body.

This limitation is particularly significant given that aging is not a uniform process across all biological systems. Different functional modules within an organism may age at varying rates, influenced by factors such as environmental exposures, genetic differences, and specific interventions. For instance, calorie restriction (CR) has been shown to primarily affect glucose metabolism pathways, while rapamycin treatment predominantly impacts the mTOR pathway ^14^. Recent studies on transcriptomics provide further evidence for this heterogeneity in aging processes. For example, cancer has been associated with pro-aging inflammatory responses and anti-aging changes in differentiation and ECM organization modules, while Klotho knock-out models exhibit accelerated aging in respiration and energy metabolism pathways, but not in inflammation ^15^. These observations underscore the need for a more mechanistic approach to measuring biological age that can capture the differential aging rates across various functional modules.

To address this gap, we developed Ageome, an interpretable aging clock framework designed to simultaneously measure the epigenetic age of thousands of molecular pathways in both mice and humans. Our approach is premised on the hypothesis that the overall biological age of an organism is determined by the collective biological ages of its constituent functional modules. The age-related epigenetic changes on module-related genes may either directly impact the gene expression or reflect regulatory changes (or lack thereof) on specific functional pathways during aging. Unlike conventional epigenetic clocks, Ageome provides a distribution of biological ages across different functional modules, offering a more comprehensive and granular view of the aging process. By applying Ageome to various models of aging and longevity interventions, we aim to establish a deeper understanding of rejuvenation and age acceleration mechanisms. This approach not only allows for the identification of pathways most affected by specific interventions but also holds the potential to inform the development of targeted anti-aging strategies through a high-dimensional representation of biological aging.

## Results

### Constructing Ageome clocks

We first obtained DNA methylation profiles of whole blood from 141 mice (C57Bl/6, 3- to 35-month-old, 16 age groups) ^16^, and 2,664 human subjects ^17,18^. Biological features were assigned to CpG sites based on the annotations of nearby genes and cis-regulatory regions (Figure 1a, Methods) ^19^. Combining pathways and epigenetic information and applying elastic net regression, we constructed aging clocks for each pathway from Kyoto Encyclopedia of Genes and Genomes (KEGG), Reactome, and Hallmark geneset ^20–22^, resulting in 1,863 Ageome clocks.

**Figure 1.**
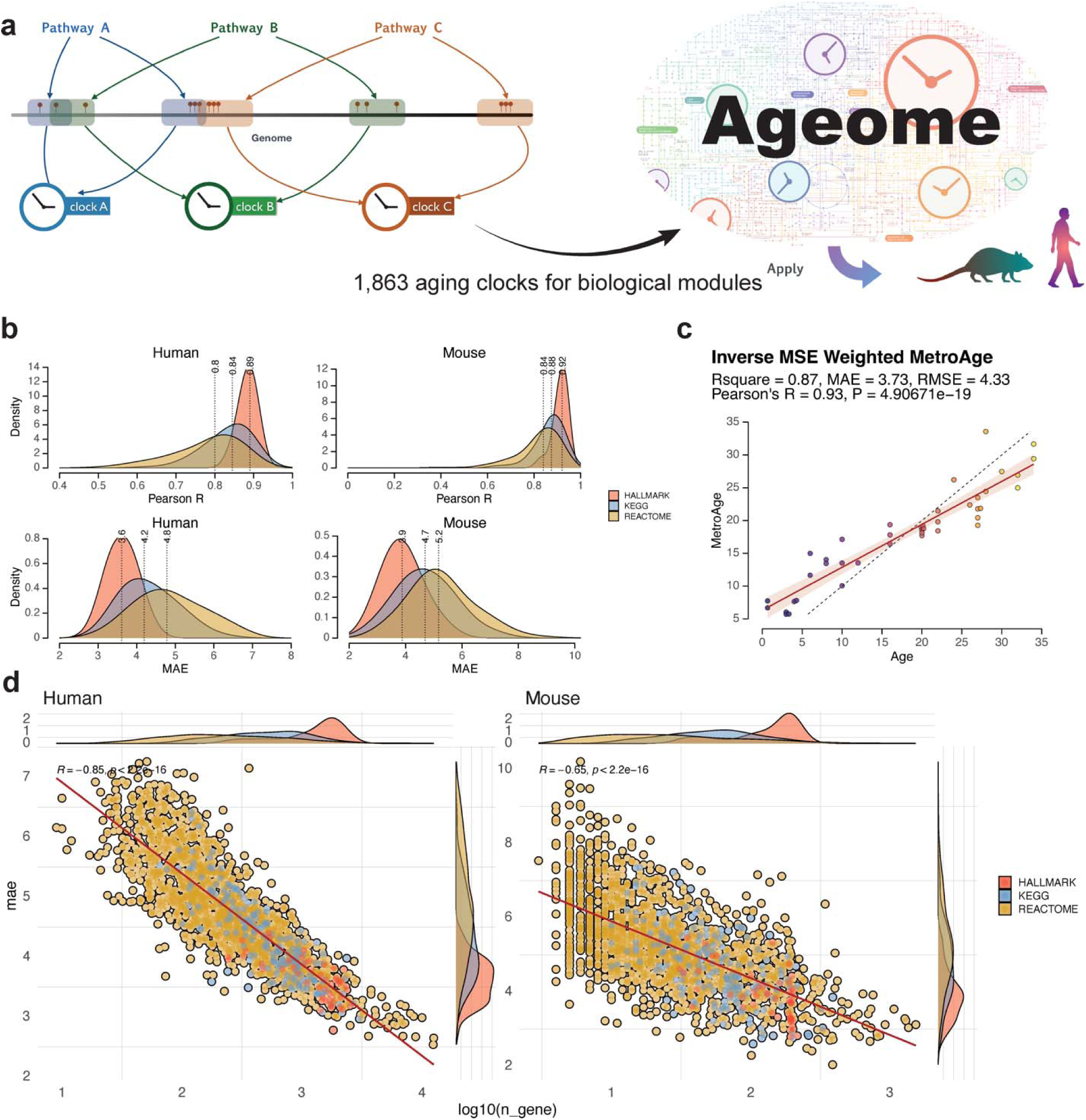
Ageome clock offers epigenetic age estimates for functional modules. (**a**) Schematic plot showing the workflow of the Ageome clock. (**b**) Density plots showing the distribution of Pearson’s R and mean absolute error (MAE) for each Ageome clock (Hallmark, KEGG, Reactome) in the test set. The results show that Ageome clocks for most pathways accurately predict the chronological age of healthy samples in both humans (left) and mice (right). (**c**) Ageome clocks are summarized and weighted by inverse root mean squared error (RMSE) in cross-validation to provide a single accurate estimate of overall biological age predicted based on all Ageome clock models in mice (MetroAge, Y-axis). The chronological age is shown on the X-axis in the unit of the month. The accuracy metrics in the test set are shown in the text. (**d**) Scatter plot showing the correlation between Ageome clock accuracy (MAE) and gene set size (at log-scale) for both human and mouse. Pearson’s R and P-values are shown in the plots. Density plots for each variable are shown in marginal plots.

In human samples, applying Ageome to the Hallmark gene set, comprising 50 pathways, resulted in a mean absolute error (MAE) of 3.70 years and Pearson’s R of 0.879 between predicted and actual age in the test set (Figure 1b). For the KEGG gene set, encompassing 186 pathways, the mean MAE was 4.53 years, and Pearson’s R was 0.812. In Reactome, the largest gene set analyzed consisted of 1600 pathways; the mean MAE was 5.12 years, accompanied by a mean Pearson’s R of 0.750. Despite the complexity and variation in different gene sets, Ageome was able to maintain a reasonable level of accuracy in predicting chronological age.

Similarly, in mouse samples, the Hallmark gene set yielded a mean MAE of 3.90 months and Pearson’s R of 0.912 (Figure 1b); KEGG pathways featured a mean MAE is 4.71 months and Pearson’s R of 0.866; and Reactome pathways exhibited a mean MAE of 5.28 months and Pearson’s R of 0.818. These results collectively demonstrate the consistent performance of the Ageome framework across various functional modules, suggesting its utility in studying aging mechanisms across species. To obtain a single summary of Ageome, we also calculated the inverse-error-weighted average of all components of Ageome (termed as MetroAge, Figure 1c). MetroAge provided an accurate estimate of the age of the mice, with an MAE of 3.73 months and a Pearson’s R of 0.93.

### Different Ageome modules show distinct predictive power of aging

We hypothesized that the difference in performance across different biological pathways is driven by two factors: 1) the number of genes in the gene set, and 2) the association of epigenetic changes in gene regulatory regions with age. To test this hypothesis, we performed a regression analysis between the MAE and the number of genes in the gene set (Figure 1d). There was a significant inverse linear relationship between test MAE and log-transformed number of genes in the gene set for both human (p < 2.2e-16, Pearson’s R = -0.85) and mouse (p < 2.2e-16, Pearson’s R = -0.65). These results suggest that the performance of Ageome clocks is partially driven by the number of genes in the gene set. To understand the association between Ageome and aging, we calculated the adjusted MAE for each pathway by regressing out the number of genes in the pathway. Upon adjusting for the number of genes in pathways, we found that the pathways most predictive of aging in humans were primarily associated with lipid metabolism (Figure 2a). The top five lipid metabolism pathways were: Linoleic acid metabolism (residual -1.51), Synthesis of very long chain fatty acyl CoAs (residual -1.27), Alpha-linolenic (omega 3) and linoleic (omega 6) acid metabolism (residual -1.16), Biosynthesis of unsaturated fatty acids (residual - 1.13), and Fatty acyl CoA biosynthesis (residual -1.04). Conversely, pathways that were least predictive of aging (those with positive residuals) were mainly related to immune system regulation and DNA replication. These include the Activation of C3 and C5 (residual 3.11), Assembly of the ORC complex at the origin of replication (residual 2.64), and E2F-enabled inhibition of pre-replication complex formation (residual 2.34).

**Figure 2.**
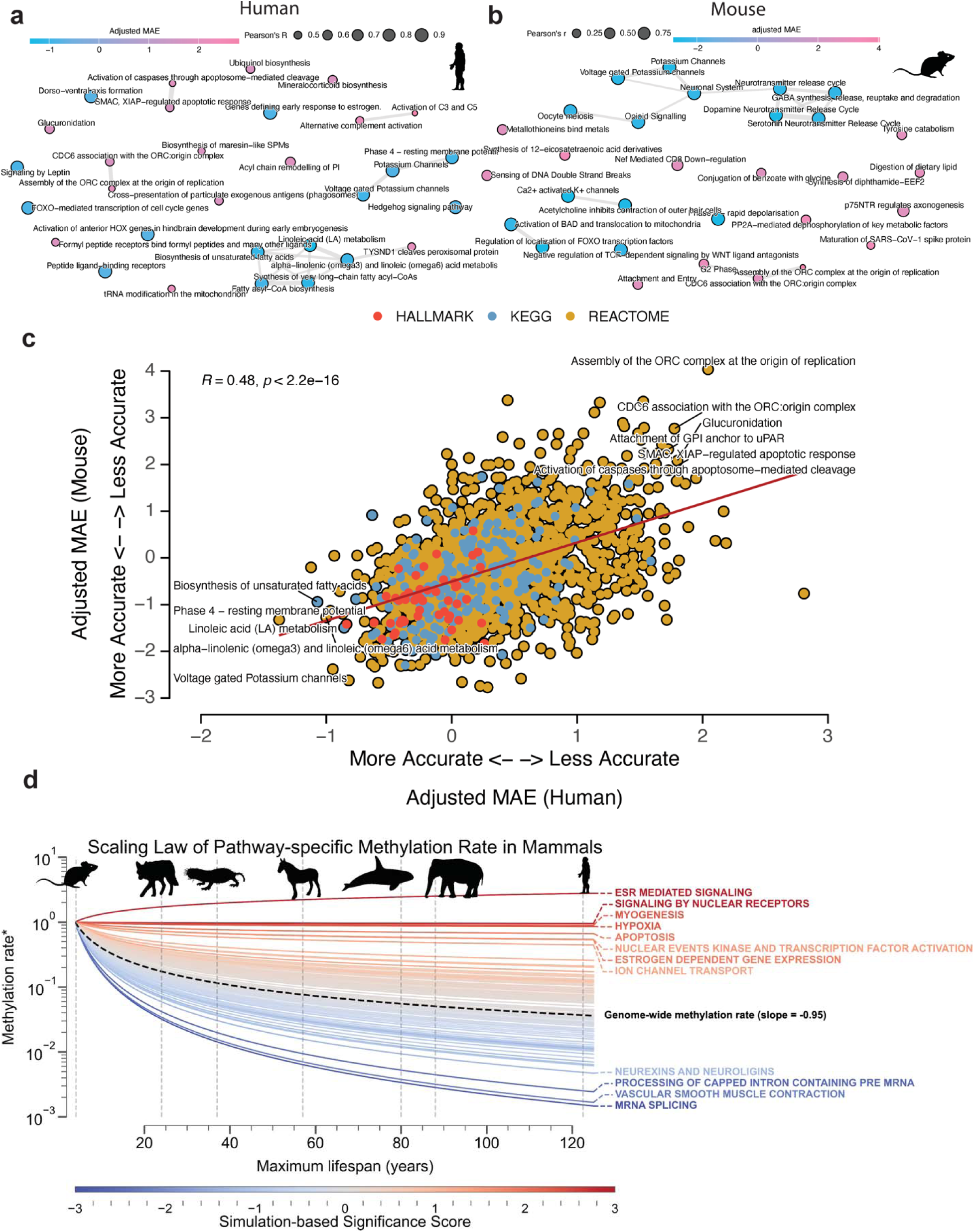
Different pathways vary in their associations with aging. (**a**, **b**) The most accurate and inaccurate Ageome clock pathways after adjusting for the size of gene sets for human (**a**) and mouse (**b**). The circle color shows adjusted mean absolute error (MAE), and the size shows Pearson’s R in the test set. Pathways with overlapped genes are clustered together with connected bonds. (**c**) Scatter plot showing the correlation between adjusted Ageome clock accuracy (MAE) in humans and mice. Pearson’s R and P-values are shown in the plot. The top accurate (left bottom) and inaccurate (right top) Ageome clocks are annotated. (**d**) Scaling law analysis of pathway-specific methylation rates across mammalian species with different maximum lifespans. Each line represents a specific pathway, color represents the -log10 P-value based on the simulation, with red lines indicating pathways that maintain relatively stable methylation rates across species lifespans, and blue lines showing pathways with more rapid declines in methylation rates as lifespan increases. The dashed black line represents the genomewide average methylation rate scaling. The x-axis shows the maximum lifespan of different mammalian species (representative species are annotated with gray dashed line and silhouettes), while the y-axis displays the methylation rate (*ratio compared to baseline species) on a logarithmic scale. Pathways with nominal P-value < 0.05 are annotated.

Similarly, we found that the top age-predicting pathways in mice were predominantly associated with ion transport and neurological signaling (Figure 2b). The top five were: Ca2+ activated K+ channels (residual -2.81), Regulation of localization of FOXO transcription factors (residual - 2.79), Potassium channels (residual -2.76), Phase 0 rapid depolarisation (residual -2.75), and Dopamine neurotransmitter release cycle (residual -2.75). In contrast, pathways that were least predictive of aging were mainly related to DNA replication and various metabolic processes. These include the Assembly of the ORC complex at the origin of replication (residual 4.01), Synthesis of 12-eicosatetraenoic acid derivatives (residual 3.35), and Maturation of SARS-CoV-1 spike protein (residual 3.33).

We also found that the adjusted MAE across pathways is significantly conserved between humans and mice (Pearson’s R = 0.51, p-value < 2.2e-16, Figure 2c). Notably, pathways related to voltage-gated potassium channels, hedgehog signaling, and genes defining early response to estrogen exhibited strong negative residuals in both species. Similarly, lipid metabolism pathways, including linoleic acid metabolism, alpha-linolenic (omega3) and linoleic (omega6) acid metabolism, and biosynthesis of unsaturated fatty acids, emerged as highly accurate age predictors. On the other hand, pathways that showed less predictive power and hence had positive residuals were primarily related to DNA replication and immune system regulation. These include E2F-enabled inhibition of pre-replication complex formation, CD22-mediated BCR regulation, attachment of GPI anchor to uPAR, CDC6 association with the ORC: origin complex, and assembly of the ORC complex at the origin of replication. In summary, these findings underscore the potential of certain conserved pathways to accurately predict age across different species. This suggests that, despite complexity of aging, there are common biological underpinnings that are reflected in these conserved pathways.

To further investigate the evolutionary conservation of age-related epigenetic changes across species, we performed a scaling law analysis of pathway-specific methylation rates in 42 mammals with varying maximum lifespans (Figure 2d). We observed a general trend of decreasing methylation rates with increasing maximum lifespan across mammalian species, consistent with previous findings of slower epigenetic aging in longer-lived species ^23^. However, the rate of this decrease varied substantially among different pathways. Upon further investigation, we found that while the average scaling law was independent of the pathway size, our confidence in the inference decreased with the size of the pathway (Extended Data Figure S1 a, b). For this purpose, we developed a size-sensitive null hypothesis based on random simulations of pathways of different sizes and associated a two-tailed p-value to each pathway (Extended Data Figure S1 c). These results suggest that evolutionary pressures on longevity affect various biological processes proportionally to the age-related number of sites included in pathways.

Notably, twelve pathways exhibited significantly different methylation rates scaling patterns across species lifespans compared to the genome-wide baseline. Three of them remain significant after being corrected for multiple tests with FDR, namely ESR-mediated signaling, signaling by nuclear receptors, and mRNA splicing. Among them, ESR-mediated signaling and signaling by nuclear receptors show a significantly slower or no decline in methylation rates with increasing lifespan, suggesting that these pathways may play conserved roles in aging processes across mammals, regardless of species-specific longevity. In contrast, mRNA splicing showed more rapid declines in methylation rates with increasing lifespan, hinting at potential adaptive changes in longer-lived species, possibly reflecting more efficient regulation of this process in animals with extended lifespans.

### Ageome predicts mortality risk and reveals hallmark agers

To assess the clinical relevance of Ageome, we applied our framework to the Normative Aging Study (NAS) cohort, comprising 1,488 individuals with a 38.8% mortality rate (Figure 3a). We analyzed the association between 1,863 Ageome clocks and mortality risk, revealing significant heterogeneity in the predictive power of different pathways. Among the clocks tested, 1,506 Ageome clocks showed a significant association with mortality risk after correction for multiple testing (FDR < 0.05, Figure 3a). Several pathways, including gluconeogenesis, RHOF GTPase cycle, and amino acid regulation of mTORC1, exhibited both high statistical significance and hazard ratios for mortality risk. Interestingly, there are 11 models that show inverse association with mortality, similar to AdaptAge, suggesting that these modules may potentially contribute to protective changes during aging. We also showed that the accuracy of Ageome clocks was only weakly correlated with their predictive power for mortality (Figure 3b). This suggests that the biological processes most indicative of chronological age may not necessarily be the most predictive of mortality risk.

**Figure 3.**
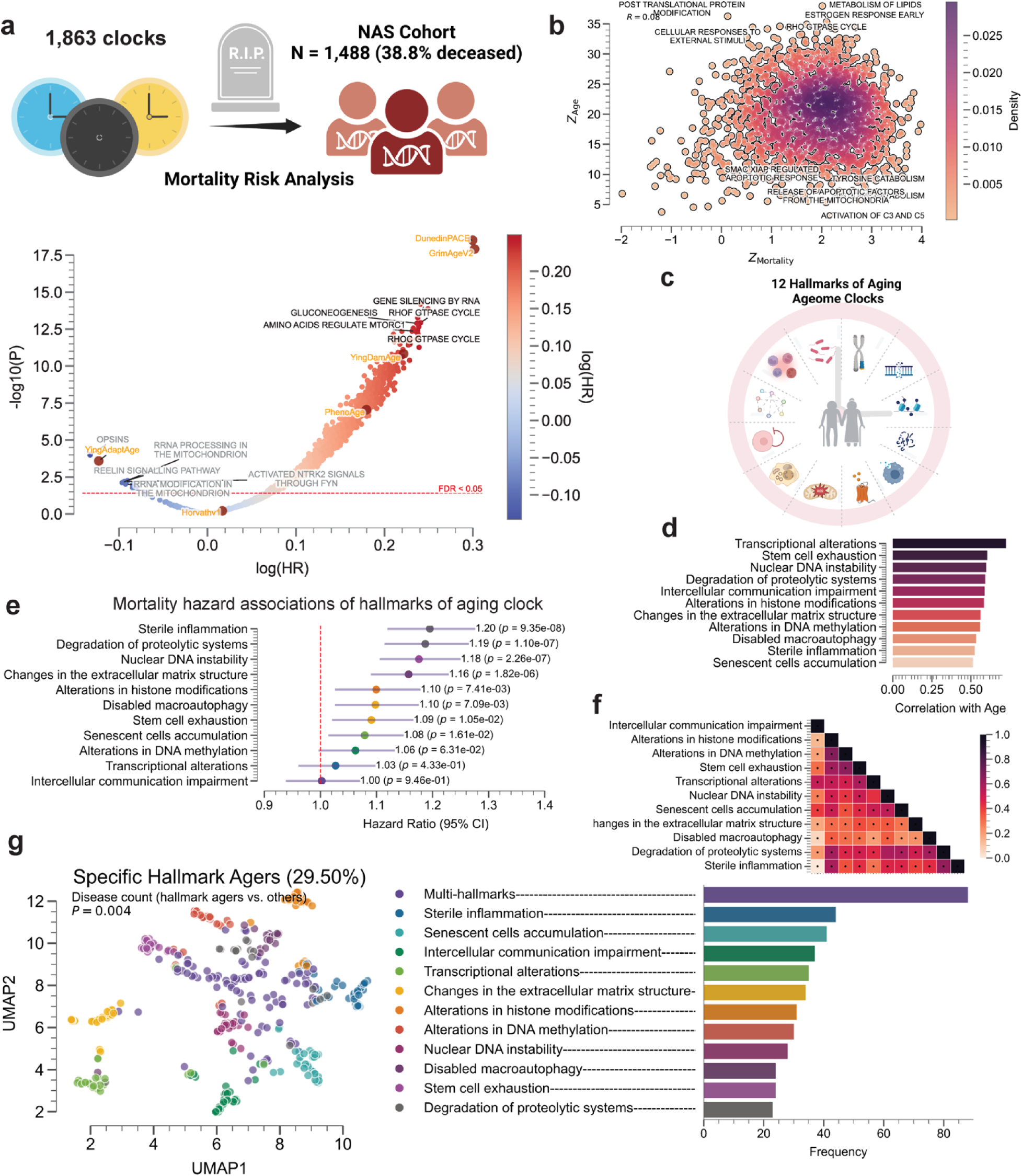
Ageome clock predicts mortality risk and reveals hallmark agers. (**a**) Mortality risk analysis of 1,863 Ageome clocks in the NAS cohort (N = 1,488, 38.8% deceased). Volcano plot showing the relationship between log(HR) and -log10(Adjusted-P) for different pathways. Top 5 pathways with high significance and hazard ratios, and with R-square > 0.5 in age prediction are labeled in black. The top 5 pathways with a negative association with mortality are labeled in gray. The red dashed line shows the FDR threshold of 0.05. (**b**) Density plot showing the relationship between Z-scores for mortality risk prediction and Ageome clock accuracy. Notable pathways are labeled. Pearson’s correlation coefficient is annotated at the left top corner of the plot in text. (**c**) Schematic representation of the 12 Hallmarks of Aging Ageome clocks analyzed. (**d**) Bar plot showing the correlation of different hallmarks of aging Ageome clocks with chronological age. (**e**) Forest plot displaying mortality risk (hazard ratios with 95% CI) for Hallmarks of Aging Ageome clocks. P-values are shown for each hallmark. (**f**) Heatmap showing the correlation between age deviation terms of different hallmarks of aging Ageome clocks, colored by Pearson’s R. The significant correlations after correcting for multiple testing using Bonferroni correction are annotated with black dots. (**g**) UMAP plot visualizing specific hallmark agers (29.50% of the cohort). Each point represents an individual, colored by their dominant hallmark of aging. To cluster, we identified individuals with any hallmark age deviation Z-score > 1.5, and converted all Z-score smaller than this threshold to 0. Individuals with more than 2 hallmark age deviations larger than 1.5 are annotated as multi-hallmark agers. Bar chart showing the frequency of different hallmark agers in the population.

We then focused our analysis on the 12 hallmarks of aging, which have Pearson’s correlation coefficient of at least 0.5 with chronological age (Figure 3c-d, Methods). Notably, transcriptional alterations, stem cell exhaustion, and nuclear DNA instability demonstrated the strongest correlations with age. When examining the relationship between hallmark clocks and mortality risk, we found that sterile inflammation, degradation of proteolytic systems, and nuclear DNA instability were associated with the highest hazard ratios (Figure 3e). These particular hallmarks may therefore play a more significant role in determining lifespan. Moreover, the age deviation term of all 12 hallmarks shows a significant positive correlation with each other, except between nuclear DNA instability and transcriptional alterations, as well as degradation of proteolytic systems (Figure 3f).

Intriguingly, our clustering analysis identified distinct clusters of individuals who show positive age deviation on specific hallmark pathways (i.e., specific hallmark agers, Figure 3g). These individuals, representing 29.50% of the cohort, showed accelerated aging primarily in one or a few specific hallmarks. The distribution of these hallmark agers varied, with multi-hallmark agers being the most common, followed by those predominantly affected by sterile inflammation and senescent cell accumulation (Figure 3g). These hallmark agers tended to have more prior health conditions compared to non-hallmark agers based on regression analysis on disease count (P = 0.004). This finding suggests that aging trajectories assessed at the epigenetic level can differ significantly between individuals, consistent with previous reports leveraging alternative omics measurements ^24–26^. Some people may experience accelerated aging in specific biological processes while maintaining relative youth in others. This heterogeneity in aging patterns could have important implications for personalized approaches to age-related interventions and preventive strategies.

### Ageome predicts the risk of diseases and separates disease types

To evaluate clinical utility of Ageome, we applied our framework to the Mass General Brigham (MGB) cohort, comprising 4,246 individuals (Figure 4a) ^27^. We assessed the association between 1,863 Ageome clocks and the risk of 43 diseases (including three general disease categories), spanning cardiovascular, cancer, respiratory, liver, and other conditions, comparing with four state-of-the-art published models (GrimAgeV2, DunedinPACE, PhenoAge, and YingDamAge) based on previous benchmarking result (Table 1)^28^.

**Figure 4.**
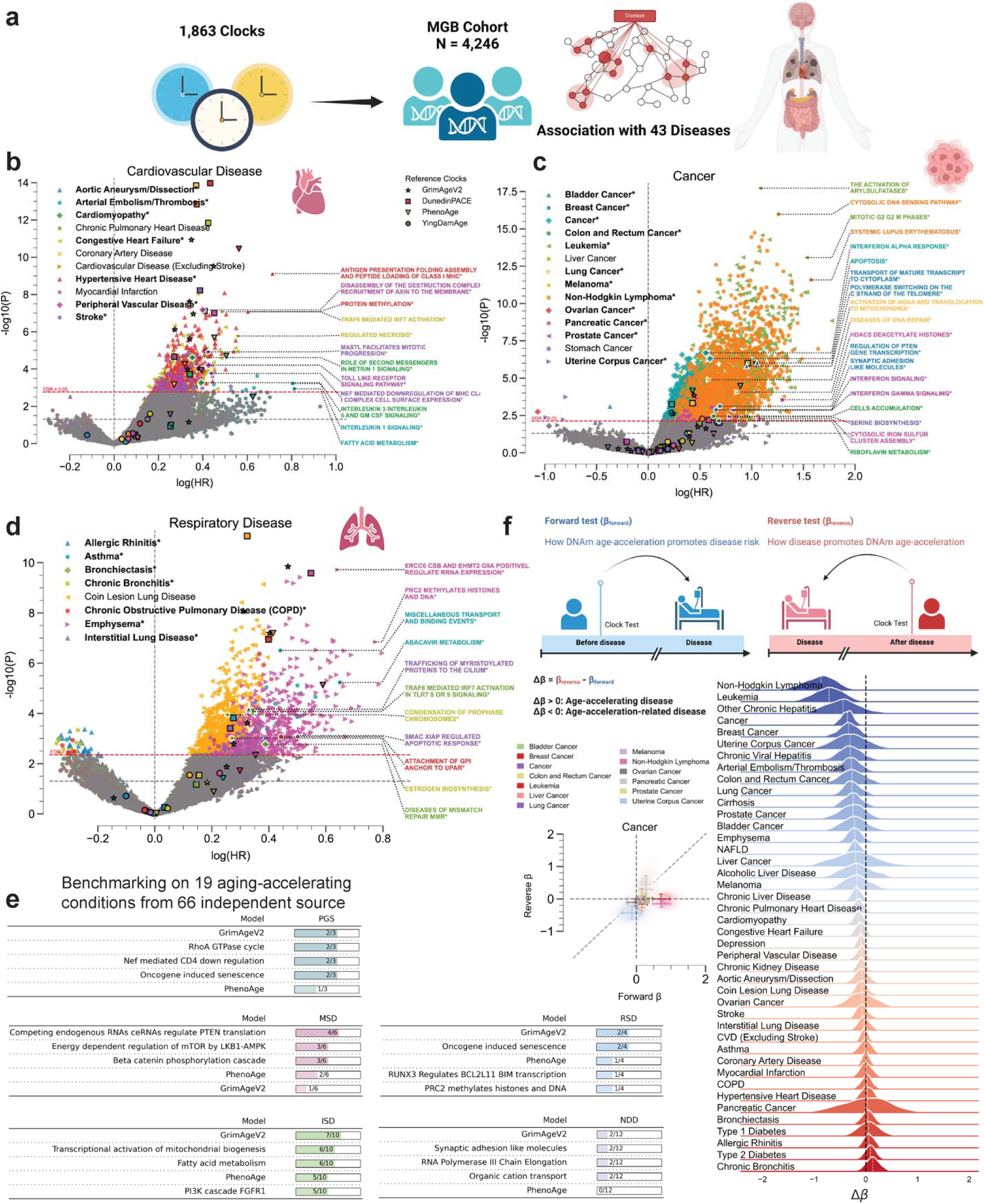
Ageome predicts disease risk and reveals bidirectional aging-disease relationships. (**a**) Schematic of the Ageome clock framework applied to the Mass General Brigham (MGB) cohort (N = 4,245) for disease risk prediction. (**b**-**d**) Volcano plots showing the relationship between log(HR) and -log10(P) for Ageome clock predictions of (**b**) cardiovascular diseases, (**c**) cancers, and (**d**) respiratory diseases. Top two pathways for disease with FDR < 0.05 are labeled. Reference models are indicated by distinct markers with black borders: GrimAgeV2 (star), DunedinPACE (square), PhenoAge (triangle), and YingDamAge (circle). FDR threshold is set to 0.05, indicated by the dashed red line. The diseases that have Ageome measurements that outperform (either by significance or hazard ratio) reference models are bolded and indicated by an asterisk (*), as well as the outperforming pathways. Only the top 2 Ageome pathways (ranked by -log10(P) * log(HR)) with FDR < 0.05 are shown for each disease. (**e**) Benchmarking analysis comparing Ageome clocks to GrimAgeV2 and PhenoAge across various disease categories using the ComputAgeBench framework. ISD: immune system diseases; MSD: musculoskeletal diseases; NDD: neurodegenerative diseases; PGS: progeroid syndromes; RSD: respiratory diseases. Panels for cardiovascular diseases and metabolic diseases are not shown as none of the clocks provide predictions for these conditions. (**f**) Bidirectional analysis of aging-disease relationships. Schematic plot showing the logistics of forward (βForward) and reverse (βReverse) analysis. Scatter plot and 2D density plot (left) shows forward (X-axis) and reverse (Y-axis) effects for different cancers. Error bar shows the standard deviation of the forward and reverse effects across Ageome clock measurements. Delta age (Δβ) represents the difference between forward and reverse effects. The density plots show the distribution of Ageome clock Δβ for each disease, with the white dashed line annotating the median value. The black dashed line shows the line where Δβ = 0.

**Table 1.** Many Ageome clocks outperform established clocks in predicting age-related diseases.

| Disease | Best Ageome Clock | P | HR | Best Reference Clock | P | HR |
| --- | --- | --- | --- | --- | --- | --- |
| Alcoholic Liver Disease | Transport of Nucleotide Sugars | 2.93e-04 | 2.63 | YingDamAge | 2.06e-02 | 1.72 |
| Allergic Rhinitis | SLBP Dependent Processing of Replication Dependent Histone Pre mRNAs | 2.17e-02 | 1.35 | DunedinPACE | 5.54e-01 | 1.03 |
| Aortic Aneurysm/Dissection | Polo Like Kinase Mediated Events | 4.00e-03 | 1.26 | DunedinPACE | 3.29e-01 | 1.10 |
| Arterial Embolism/Thrombosis | Interleukin 1 Signaling | 5.96e-04 | 2.24 | PhenoAge | 3.15e-03 | 1.87 |
| Asthma | Abacavir Metabolism | 5.82e-06 | 1.91 | DunedinPACE | 1.52e-04 | 1.32 |
| Bladder Cancer | Transport of Mature Transcript to Cytoplasm | 9.61e-07 | 2.86 | PhenoAge | 9.04e-03 | 1.92 |
| Breast Cancer | Regulation of PTEN Gene Transcription | 7.65e-04 | 2.00 | PhenoAge | 2.17e-01 | 1.36 |
| Bronchiectasis | TRAF6 Mediated IRF7 Activation in TLR7 8 or 9 Signaling | 8.34e-05 | 1.41 | GrimAgeV2 | 1.34e-02 | 1.29 |
| Cancer | Apoptosis | 4.90e-07 | 1.82 | PhenoAge | 7.11e-05 | 1.53 |
| Cardiomyopathy | Role of Second Messengers in Netrin 1 Signaling | 2.42e-05 | 1.42 | GrimAgeV2 | 9.52e-05 | 1.47 |
| Chronic Bronchitis | Condensation of Prophase Chromosomes | 1.14e-04 | 1.29 | DunedinPACE | 2.91e-02 | 1.17 |
| Chronic Liver Disease | MASTL Facilitates Mitotic Progression | 1.46e-06 | 1.54 | DunedinPACE | 7.98e-12 | 1.48 |
| Chronic Viral Hepatitis | O Glycan Biosynthesis | 1.60e-03 | 2.05 | PhenoAge | 1.04e-01 | 1.69 |
| Cirrhosis | Mitotic G1 Phase and G1 S Transition | 7.47e-05 | 1.79 | DunedinPACE | 4.93e-07 | 1.63 |
| Coin Lesion Lung Disease | SEMA4D Mediated Inhibition of Cell Attachment and Migration | 7.24e-10 | 1.46 | DunedinPACE | 8.63e-12 | 1.38 |
| Colon and Rectum Cancer | Cells Accumulation | 3.53e-03 | 1.88 | GrimAgeV2 | 3.22e-01 | 1.25 |
| Congestive Heart Failure | TRAF6 Mediated IRF7 Activation | 8.48e-08 | 1.62 | DunedinPACE | 1.44e-12 | 1.53 |
| Emphysema | ERCC6 CSB and EHMT2 G9A Positively Regulate rRNA Expression | 1.90e-10 | 1.90 | DunedinPACE | 2.59e-10 | 1.73 |
| Hypertensive Heart Disease | Antigen Presentation Folding Assembly and Peptide Loading of Class I MHC | 7.61e-10 | 2.04 | DunedinPACE | 1.07e-14 | 1.54 |
| Interstitial Lung Disease | SMAC XIAP Regulated Apoptotic Response | 7.64e-04 | 1.65 | GrimAgeV2 | 2.42e-04 | 1.38 |
| Leukemia | Mitotic G2 G2 M Phases | 8.35e-14 | 4.61 | PhenoAge | 3.39e-05 | 2.44 |
| Lung Cancer | Diseases of DNA Repair | 8.80e-05 | 2.37 | GrimAgeV2 | 1.43e-04 | 1.56 |
| Melanoma | FGFR2 Alternative Splicing | 1.77e-02 | 1.80 | PhenoAge | 4.03e-01 | 1.28 |
| Non-Hodgkin Lymphoma | Cytosolic DNA Sensing Pathway | 1.06e-16 | 3.52 | PhenoAge | 1.21e-06 | 2.83 |
| Other Chronic Hepatitis | Degradation of Cysteine and Homocysteine | 1.65e-02 | 5.00 | PhenoAge | 5.64e-02 | 2.16 |
| Ovarian Cancer | Cargo Trafficking to the Periciliary Membrane | 4.78e-02 | 2.19 | PhenoAge | 5.74e-01 | 1.35 |
| Pancreatic Cancer | HDACs Deacetylate Histones | 2.83e-04 | 3.65 | DunedinPACE | 6.32e-03 | 1.80 |
| Peripheral Vascular Disease | Disassembly of the Destruction Complex and Recruitment of AXIN to the Membrane | 6.65e-08 | 1.49 | GrimAgeV2 | 2.48e-06 | 1.41 |
| Prostate Cancer | Interferon Signaling | 1.38e-03 | 2.25 | PhenoAge | 6.82e-01 | 1.12 |
| Stomach Cancer | Assembly of the HIV Virion | 5.85e-02 | 1.60 | YingDamAge | 7.37e-01 | 1.12 |
| Stroke | MASTL Facilitates Mitotic Progression | 1.21e-05 | 1.53 | DunedinPACE | 6.02e-09 | 1.47 |
| Type 1 Diabetes | The NLRP3 Inflammasome | 1.78e-07 | 1.79 | DunedinPACE | 1.06e-05 | 1.81 |
| Uterine Corpus Cancer | Serine Biosynthesis | 3.91e-03 | 2.21 | DunedinPACE | 9.66e-03 | 1.99 |
| Cardiovascular Disease (Excluding Stroke) | CD28 Dependent PI3K AKT Signaling | 1.93e-04 | 1.32 | DunedinPACE | 1.44e-13 | 1.45 |
| Chronic Kidney Disease | IRAK1 Recruits IKK Complex | 5.80e-08 | 1.36 | DunedinPACE | 1.52e-11 | 1.41 |
| Chronic Obstructive Pulmonary Disease (COPD) | LRR FLII Interacting Protein 1 LRRFIP1 Activates Type I IFN Production | 4.28e-07 | 1.44 | GrimAgeV2 | 6.46e-08 | 1.50 |
| Chronic Pulmonary Heart Disease | Translesion Synthesis by POLK | 1.60e-06 | 1.40 | GrimAgeV2 | 3.09e-10 | 1.57 |
| Coronary Artery Disease | RIPK1 Mediated Regulated Necrosis | 6.62e-04 | 1.33 | DunedinPACE | 1.42e-14 | 1.45 |
| Depression | Caspase Activation via Death Receptors in the Presence of Ligand | 1.61e-04 | 1.34 | DunedinPACE | 1.76e-08 | 1.36 |
| Liver Cancer | Degradation of Beta Catenin by the Destruction Complex | 3.03e-04 | 2.19 | DunedinPACE | 9.80e-07 | 2.62 |
| Myocardial Infarction | IRAK1 Recruits IKK Complex | 3.00e-03 | 1.33 | DunedinPACE | 1.00e-07 | 1.57 |
| Non-Alcoholic Fatty Liver Disease (NAFLD) | Regulation by C FLIP | 1.59e-05 | 1.39 | DunedinPACE | 8.83e-09 | 1.41 |
| Type 2 Diabetes | STAT5 Activation | 7.94e-06 | 1.42 | DunedinPACE | 5.56e-27 | 1.89 |

The table shows the top Ageome clock and the top established clock for predicting various age-related diseases. The P-value and hazard ratio (HR) are shown for both clocks. Ranking of the clocks are based on $-\log_{10}(\text{P-value}) * \log(\text{HR})$ .

For cardiovascular diseases (Figure 4b), Ageome showed superior performance for 6 out of 11 conditions. Notably, for arterial embolism/thrombosis, the Interleukin 1 signaling pathway (HR = 2.24, p = 5.96e-04) and fatty acid metabolism (HR = 2.25, p = 1.16e-03) were particularly predictive. In cardiomyopathy, the Role of second messengers in netrin 1 signaling pathway (HR = 1.42, p = 2.42e-05) outperformed existing models. For peripheral vascular disease, the Disassembly of the destruction complex and recruitment of axin to the membrane pathway (HR = 1.49, p = 6.65e-08) showed superior predictive power.

In cancer prediction (Figure 4c), Ageome outperformed traditional risk factors for multiple cancer types (8 out of 14, including general cancer, which includes all cancer subtypes). For bladder cancer, the Transport of mature transcript to cytoplasm pathway (HR = 2.86, p = 9.61e-07) and Polymerase switching on the C strand of the telomere pathway (HR = 2.59, p = 1.58e-06) were highly predictive. In leukemia, the Mitotic G2 G2 M phases pathway (HR = 4.61, p = 8.35e-14) showed remarkable predictive power. For lung cancer, the Diseases of DNA repair pathway (HR = 2.37, p = 8.80e-05) outperformed existing models. Non-Hodgkin lymphoma (NHL) prediction was significantly improved by the Cytosolic DNA sensing pathway (HR = 3.52, p = 1.06e-16), as well as Systemic Lupus Erythematosus (SLE) pathway (HR = 4.83, p = 2.64e-12), consistent with clinical observations that the SLE patients are at greater risk for NHL ^29^.

For respiratory diseases (Figure 4d), Ageome again demonstrated superior predictive capabilities (7 out of 8), particularly for asthma and bronchiectasis. The Miscellaneous transport and binding events pathway (HR = 1.55, p = 3.13e-07) was highly predictive for asthma, while the TRAF6 mediated IRF7 activation in TLR7 8 or 9 signaling pathway (HR = 1.41, p = 8.34e-05) showed superior performance for bronchiectasis. In liver diseases (Figure S2), Ageome still provided valuable insights. For instance, the MASTL facilitate mitotic progression pathway was highly associated with non-alcoholic liver disease and general chronic liver disease risk. For other diseases, Ageome showed particular strength in predicting Type 1 Diabetes, with the NLRP3 inflammasome pathway (HR = 1.79, p = 1.78e-07) and DARPP 32 events pathway (HR = 1.65, p = 3.80e-08) outperforming existing predictors, which agrees with the investigated relationship between NLRP3 inflammasome and T1D ^30^.

To further validate our findings, we conducted an independent benchmarking analysis (Figure 4e) comparing Ageome to established aging clocks, specifically GrimAgeV2 and PhenoAge, using the ComputAgeBench framework ^31^. This analysis evaluated the ability of these clocks to differentiate between aging acceleration conditions and healthy control samples across various disease categories, namely immune system diseases (ISD), musculoskeletal diseases (MSD), neurodegenerative diseases (NDD), progeroid syndromes (PGS), and respiratory diseases (RSD), metabolic diseases, and cardiovascular diseases. Ageome consistently matched the predictive power of GrimAgeV2 and PhenoAge across the examined disease categories.

To investigate the bidirectional relationship between disease onset and epigenetic age acceleration, we employed a novel bidirectional analysis approach using the MGB dataset. We conducted two types of tests: forward (βForward) and reverse (βReverse) for each of the Ageome predictors. The forward test assessed how DNAm age acceleration might promote disease risk, while the reverse test examined how disease onset could potentially accelerate epigenetic aging. Delta beta (Δβ) represents the difference between the forward (βForward) and reverse (βReverse) effects (i.e., βReverse - βForward) for each disease. It quantifies the net direction and magnitude of the relationship between epigenetic age acceleration and disease. A negative Δβ indicates a stronger tendency for accelerated aging to precede disease onset, while a positive Δβ suggests that disease onset more strongly influences subsequent epigenetic age acceleration. The distribution of this metric across Ageome predictors provides a comprehensive summary of the dominant direction in the aging-disease relationship for each condition studied.

Our analysis uncovered distinct patterns across various diseases (Figure 4f, Figure S3). Certain conditions, including various cancers, other chronic hepatitis, and arterial embolism/thrombosis, showed stronger effects in the forward direction (βForward > βReverse), indicating that accelerated epigenetic aging may be a more significant factor in their onset. Conversely, chronic bronchitis and type 2 diabetes displayed stronger effects in the reverse direction (βReverse > βForward), suggesting that their onset may have a more pronounced impact on accelerating the epigenetic aging process. Overall, this suggests that while accelerated aging can increase disease risk for many conditions, the onset of certain diseases may also contribute to further acceleration of the aging process. This complex interaction underscores the importance of considering both directions when studying age-related diseases and developing interventions.

### Ageome reveals potential functional impacts of established longevity interventions

We then applied Ageome to various models of established longevity interventions, including calorie restriction (CR), Snell dwarf mice, growth hormone receptor knockout, iPSC reprogramming, and heterochronic parabiosis (Figure 5, Figure S4) ^16,32^. Since Ageome provides the distribution of biological ages across functional modules, it offers an estimation of longevity effects with regard to individual pathways, thereby helping to identify major pathways and mechanisms associated with each of these models. We thus identified pathways that are primarily rejuvenated by each intervention, as well as common signatures of known lifespan-extending interventions (Figure 5).

**Figure 5.**
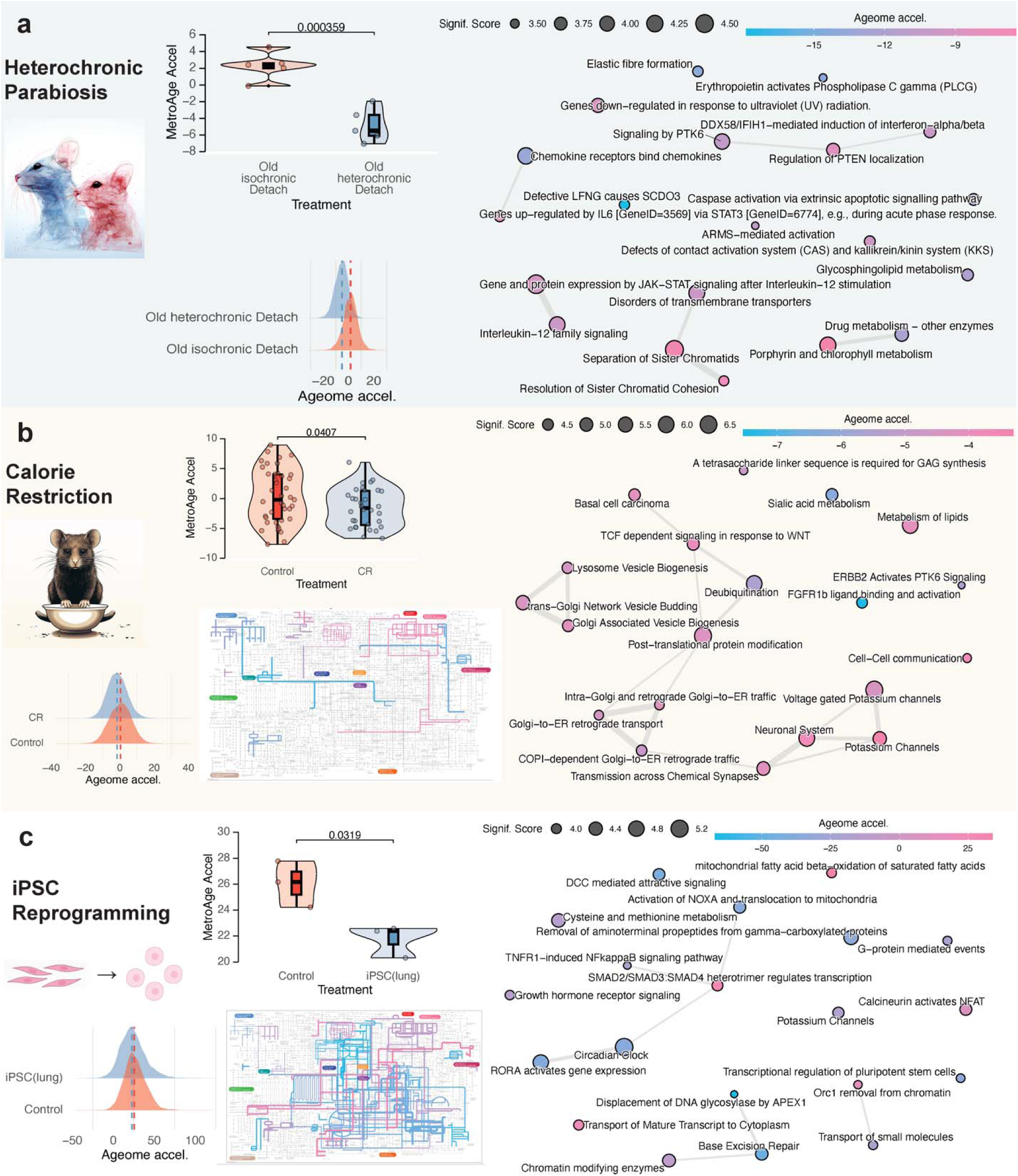
Different longevity interventions show distinct patterns in Ageome. (**a**-**c**) Application of Ageome clocks to heterochronic parabiosis model after detachment (**a**), caloric restriction (**b**), and iPSC reprogramming of lung fibroblasts (**c**). MetroAge and the distribution of Ageome clock prediction show the overall effect of interventions (left). P-values are shown in text. Network plot shows connections across affected Ageome clocks (right). Only top pathways with adjusted P < 0.05 are shown. Pathways with black circles are significant after being adjusted for multiple testing. The color of the dots shows the estimated biological age difference compared to controls, and the size shows -log10(P-value). Pathways with overlapped genes are clustered together with connected bonds. iPath plots provide an overview of the affected functional modules for each treatment. The color of the lines shows the estimated biological age difference compared to controls, and stroke shows -log10(P-value).

Compared to isochronic parabiosis, the summarizing MetroAge for whole blood samples of heterochronic parabiosis after detachment is significantly decelerated (p = 3.59E-04), and 421 Ageome clocks are significantly decelerated after adjusting for multiple testing (Figure 5a). Top significant decelerated pathways are related to immune system regulation, including JAK-STAT signaling after Interleukin-12 Stimulation (decreased by 9.15 months, p = 2.42E-05), chemokine receptors bind chemokines (decreased by 13.42 months, p = 5.94E-05), and DDX58/IFIH1-Mediated Induction of Interferon Alpha/Beta (decreased by 10.05 months, p = 2.12E-04). Similarly, the pathway related to heme metabolism, porphyrin, and chlorophyll metabolism (decreased by 5.64 months, p = 6.17E-05) is also decelerated. This is consistent with previous reports of broad rejuvenation induced by heterochronic parabiosis ^32^. A similar result is also observed in the parabiosis model before detachment (Figure S4).

For CR, the summarizing MetroAge is also significantly decelerated (p = 0.04) and 164 Ageome clocks are significantly decelerated (Fig 5B). We observed a significant deceleration in multiple facets of aging. Post-translational protein modification, a key process in protein biosynthesis and regulation, showed a notable deceleration of 4.31 months (p = 2.25E-07), as well as deubiquitination (5.45 months, p = 1.03E-06). Similarly, we observed significant age deceleration in mitochondria function-related pathways, including voltage-gated potassium channels (4.41 months, p = 2.62E-07) and potassium channels (3.30 months, p = 6.82E-06). Furthermore, the functional systems associated with trans-Golgi network vesicle budding demonstrated age deceleration of 4.41 months (p = 5.17E-06) and metabolism of lipids (4.07 months, p = 9.72E-07). Together, these findings underscore the impact of CR on a multitude of biological pathways.

Similarly, we also examined Ageome of growth hormone receptor knockout (GHRKO) and Snell dwarf mice (Figure S4). The summarizing MetroAges are significantly decelerated in both models (p= 0.0001 for GHRKO and p = 2.42E-05 for Snell dwarf), with 124 and 283 Ageome clocks significantly decelerated, respectively. For GHRKO, the top significant decelerated pathways are related to protein modification, including protein ubiquitination (deceleration of 6.05 months, p = 1.52E-05) and the post-translational protein modification pathway (deceleration of 4.01 months p = 7.20E-05). Several pathways related to cell cycle regulation also showed significant deceleration. This was observed in the G2 phase (5.49 months, p = 1.21E-04) and G2/M checkpoints (5.01 months, p = 1.45E-04). Concurrently, the pathway involved in the negative regulation of Notch4 signaling also experienced a marked slowdown (7.49 months, p = 3.25E-05). For Snell dwarfism, the top significant decelerated pathways are related to cellular structure, metabolism, and signaling. The adherens junction (7.17 months, p = 2.61E-07) and tight junction pathways (6.32 months, p = 6.63E-06) both show substantial deceleration. Similarly, Rho GTPase-related pathways, namely CDC42 GTPase cycle (5.08 months, p = 8.97E-06) and Rho GTPases activate CIT (7.74 months, p = 1.43E-05), were significantly slowed. Metabolic processes, including phospholipid metabolism (5.83 months, p = 1.01E-07) and glycolysis (5.87 months, p = 6.25E-06), also show significant deceleration. In the context of protein regulation, the SUMOylation of DNA damage response and repair proteins pathway shows an age deceleration of 6.69 months (p = 5.74E-06). These findings are largely aligned with the known effects of GHRKO and Snell dwarfism on aging.

For iPSC reprogramming, the summarizing MetroAge is significantly decelerated in reprogrammed lung fibroblasts (p = 0.032), and 316 Ageome clocks are significantly decelerated (Figure 5c). Interestingly, unlike other interventions tested where all significantly affected Ageome clocks show consistent deceleration, iPSC reprogramming also significantly accelerated the epigenetic age of 181 pathways (Figure 6a). The top decelerated pathway is the Circadian Clock pathway (37.41 months, p = 5.54E-06). Also, the pathways related to the regulation of gene expression and chromatin structure are highlighted by the RORA Activates Gene Expression pathway (40.25 months, p = 1.77E-05), Chromatin Modifying Enzymes pathway (9.41 months, p = 4.70E-05), and the Transcriptional Regulation of Pluripotent Stem Cells pathway (34.34 months, p = 1.17E-04). For the accelerated pathways, the key pathways are related to cell signaling and gene regulation, such as Calcineurin Activates NFAT (8.66 months, p = 6.27E-05),

**Figure 6.**
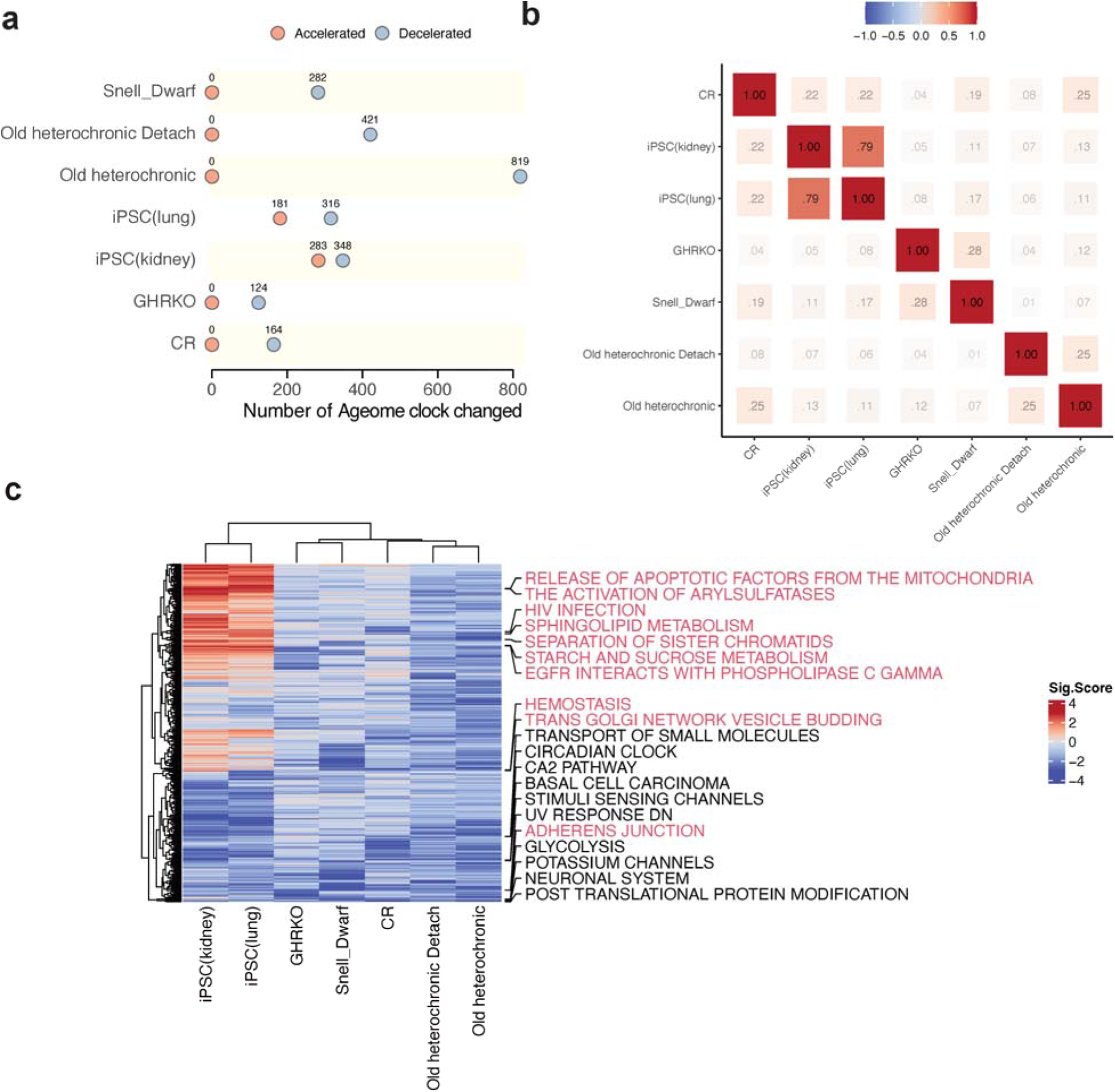
Ageome provides insights into shared and unique mechanisms of longevity interventions. (**a**) The number of Ageome clocks showing significant acceleration (red) or deceleration (blue) after adjusting for multiple testing (FDR < 0.05). (**b**) Correlation of Ageome clocks across interventions, revealing interventions with similar or different mechanisms. Spearman’s Rhos are shown in the plot. (**c**) Heatmap displaying the top 10 shared varying Ageome clocks (black), and the top 10 unique varying Ageome clocks (red) across interventions. The color indicates the signed -log10(P-value) of the Ageome clock for given interventions. The color bar is capped at 4.

SMAD2/SMAD3/SMAD4 Heterotrimer Regulates Transcription (26.78 months, p = 8.27E-05), and Transport of Mature Transcript to Cytoplasm (21.34 months, p = 9.60E-05). Metabolic pathways, like Mitochondrial Fatty Acid Beta Oxidation of Saturated Fatty Acids (33.88 months, p = 1.03E-04), exhibit accelerated aging as well. Cell cycle-related pathways like TP53 Regulates Transcription of Genes Involved in G2 Cell Cycle Arrest (23.59 months, p = 1.87E-04) and Aberrant Regulation of Mitotic Exit in Cancer due to RB1 Defects (Figure S4) also show acceleration. Similar results are also observed in reprogrammed kidney fibroblasts (Figure S4). These findings suggest that although iPSC reprogramming can rejuvenate overall epigenetic age, it may also accelerate the epigenetic age of some specific functional modules.

### Integrative analysis identifies key pathways shared by multiple interventions

Finally, we investigated the commonalities and differences across the examined interventions. We first calculated the Spearman correlation of the epigenetic age deviation predicted by the significantly affected Ageome clocks across interventions (Figure 6b). All interventions showed a generally positive correlation with each other, and three clusters emerged. The first cluster includes CR, and iPSC reprogramming for kidney and lung fibroblast; the second cluster includes GHRKO and Snell dwarfism; and the third cluster includes heterochronic parabiosis before and after detachment. Interestingly, in addition to the association with iPSCs, CR shows a relatively strong positive correlation with parabiosis before detachment and Snell dwarfism. This result suggests that CR may extend lifespan through some fundamental mechanisms shared across all examined interventions.

To comprehensively investigate the commonalities and differences across various interventions, we examined pathways implicated in the response to these interventions (Figure 6c). The top shared Ageome metrics include Transport of Small Molecules, Circadian Clock, Ca2 Pathway, Basal Cell Carcinoma, Stimuli-sensing Channels, UV Response, Glycolysis, Potassium Channels, Neuronal System, and Post Translational Protein Modification. These pathways encapsulate a diverse array of biological processes, spanning metabolism, cellular signaling, and gene regulation, indicating that they might play a significant role in the aging process and response to antiaging interventions.

Conversely, several Ageome measures exhibit considerable variation across different interventions, implying that these pathways could be specifically modulated by particular interventions. The most variable pathways include Release of Apoptotic Factors from the Mitochondria, Activation of Arylsulfatases, HIV Infection, Sphingolipid Metabolism, Separation of Sister Chromatids, Starch and Sucrose Metabolism, EGFR Interaction with Phospholipase C-gamma, Hemostasis, Trans Golgi Network Vesicle Budding, and Adherens Junction. These findings suggest that while there are some common mechanisms that are influenced by various anti-aging interventions, there are also unique pathways that are preferentially modulated by specific interventions.

## Discussion

Our study introduces the Ageome framework, a comprehensive approach to understanding biological aging at the level of DNA methylation through the lens of functional modules. This high-dimensional representation of aging offers valuable insights into the intricate mechanisms underlying this complex process, demonstrating robust performance in age prediction across human and mouse samples.

A key finding of our study is the differential predictive power of various pathways in aging. After adjusting for gene set size, we found that lipid metabolism pathways in humans and ion transport and neurological signaling pathways in mice were most predictive of aging. This aligns with previous research highlighting the central role of these processes in aging ^33,34^. The high predictability of these pathways suggests their potential as biomarkers for biological aging. Intriguingly, we observed strong cross-species conservation in aging prediction for certain pathways, particularly those related to voltage-gated potassium channels, hedgehog signaling, and early response to estrogen. This conservation points to a shared biological basis for aging across mammals ^35^. Conversely, pathways associated with DNA replication showed the least predictive power, suggesting that these fundamental cellular processes might be more stable or less predictably altered during aging.

Our analysis of the pathway-specific scaling law across different species revealed a general trend of decreasing methylation rates with increasing lifespan, consistent with previous findings ^23^. However, the variation in this trend across different pathways suggests that evolutionary pressures on longevity may have differential effects on various biological processes. This observation opens new avenues for investigating the evolutionary aspects of aging and longevity.

Application of Ageome to human cohorts yielded several significant insights into aging and disease. Our analysis of the Normative Aging Study cohort revealed that a large number of Ageome clocks (1,506 out of 1,863) were significantly associated with mortality risk. This underscores the broad impact of aging across multiple biological pathways and highlights the potential of Ageome as a comprehensive predictor of longevity. Interestingly, we found that the accuracy of Ageome clocks in predicting chronological age was only weakly correlated with their power to predict mortality risk. This suggests that the biological processes most indicative of chronological age may not necessarily be the most critical for determining lifespan. The identification of distinct “hallmark agers” - individuals showing accelerated aging primarily in specific hallmarks - suggests that aging trajectories can vary significantly between individuals, adding to a growing body of evidence that indicates that some individuals may experience accelerated aging in specific biological processes while maintaining relative youth in others ^24–26^. Further understanding of heterogeneity in aging patterns will be key for leveraging aging biomarkers to guide personalized medicine, potentially by tailoring interventions targeting the most affected hallmarks in each individual.

The application of Ageome to the Mass General Brigham cohort demonstrated its superior predictive power for a wide range of age-related diseases (27 out of 43), particularly in cancer, compared to established aging clocks (GrimAgeV2, DunedinPACE, PhenoAge, and YingDamAge). This improved predictive capability could have substantial clinical implications, potentially enabling earlier and more accurate identification of individuals at high risk for specific cancers. The consistent performance of Ageome across various disease categories in the independent ComputAgeBench framework further validates its robustness and versatility as a tool for studying aging and age-related diseases. Additionally, our bidirectional analysis, examining the relationship between disease onset and epigenetic age acceleration, provides a nuanced view of the aging-disease relationship, a key unresolved challenge in the aging field. The observation that some conditions, such as various cancers, showed stronger effects in the forward direction (epigenetic aging preceding disease onset) while others, like chronic bronchitis and type 2 diabetes, displayed stronger effects in the reverse direction (disease onset accelerating epigenetic aging) highlights the complex interplay between aging and disease. This bidirectional relationship underscores the importance of considering both preventive strategies to slow aging and interventions to mitigate the aging-accelerating effects of certain diseases.

The application of Ageome to various longevity interventions provided granular insights into their effects on different functional modules. Notably, our analysis of iPSC reprogramming revealed an unexpected picture of its impact on cellular aging. While iPSC reprogramming is generally considered to reset epigenetic age ^36–38^, our results show that this reset is not uniform across all functional modules. In fact, we observed accelerated aging of pathways following reprogramming. This finding challenges the notion of uniform rejuvenation through iPSC reprogramming and suggests a more complex process where global rejuvenation is accompanied by aging in subsets of functions.

We propose two possible explanations for this observation. First, iPSC reprogramming may promote functional modules that show protective adaptations during aging, leading to an apparent acceleration in Ageome clocks similar to the AdaptAge clock ^11^. Alternatively, the reprogramming process itself may induce cellular stress and damage, accelerating aging in certain modules. This aligns with the stochastic nature of iPSC reprogramming ^39^ and its known potential to induce cellular senescence and death ^40^. Further research is needed to elucidate whether reprogramming results in complete rejuvenation or if it comes with the side effect of accelerating some aspects of aging.

It is important to acknowledge that the relationship between DNA methylation changes and gene expression or function is complex and not always direct. While some age-related DNA methylation changes affect gene expression, other changes are not directly reflected in corresponding gene expression alterations ^41^. However, even in cases where methylation changes do not immediately affect gene expression, they may reflect upstream regulatory events, the breakdown of regulatory mechanisms, cellular responses to aging-related stressors, or alterations in the activity of transcription factors and other regulatory elements. Thus, the Ageome could provide a nuanced view of age-related changes in cellular regulation and pathway functionality, capturing both active alterations and potential regulatory deficits that may precede or accompany more overt functional changes. This perspective allows us to interpret the Ageome not just as a direct measure of pathway activity but as a sensitive barometer of age-related regulatory breakdowns across different functional modules. Future research integrating Ageome data with transcriptomic, proteomic, and functional studies will be crucial to fully elucidate the biological significance of these pathway-specific epigenetic age signatures.

Together, the Ageome framework offers a comprehensive and interpretable approach to assessing biological aging across functional modules. By expanding the traditional single numerical output of established epigenetic clocks into a rich collection of over a thousand biologically interpretable data points, Ageome demonstrates the utility of pathway-specific biological age predictions and reveals shared aging mechanisms between humans and mice. This study provides novel insights into the dynamic changes of pathway-specific epigenetic age across diverse aging interventions, including calorie restriction, iPSC reprogramming, growth hormone receptor knockout, Snell dwarfism, and heterochronic parabiosis. By highlighting both shared and unique aging mechanisms underlying these interventions. Future integration with large-scaled database (e.g., ClockBase) could potentially facilitate the development of new anti-aging strategies ^42^. Future research should focus on validating these findings in larger cohorts and across different tissues, as well as exploring the potential of Ageome-guided personalized interventions to mitigate the effects of aging.

## Methods

### Training data

The DNA methylation data for mice were obtained from the whole blood of 141 mice (C57Bl/6, 3- to 35-month-old, 16 age groups) through Reduced representation bisulfite sequencing (RRBS)^16^. The human DNA methylation data were obtained from the whole blood of 2,664 individuals through Illumina 450K array ^17,18^. Methylation data were quality-controlled and normalized using the minfi package ^43^.

### Ageome clock model

The CpG sites were annotated to genes using the rules described in GREAT ^19^. In brief, each gene is assigned a regulatory domain. This domain is composed of a basal domain that expands 5 kb upstream and 1 kb downstream from the gene’s transcription start site. Additionally, this domain extends up to the basal regulatory domain of the closest upstream and downstream genes within a 1 Mb range. All the CpG sites that occur within the domain are assigned to the gene. The CpG sites that are not assigned to any gene are excluded from the analysis.

We then further assigned the CpG sites to functional modules using various pathway databases, including KEGG, Reactome, and Hallmark ^20–22^. Hallmarks of aging pathways were collected from Open Genes ^44^. The CpG sites are included if they are annotated to the genes that are included in the pathway. All the included CpG sites are then used to train the Ageome clock model using the elastic net regression ^45^:

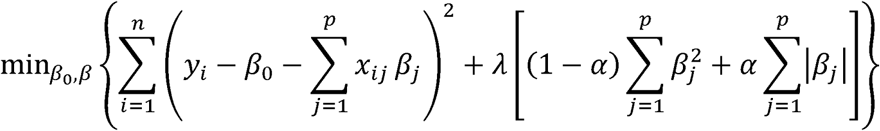

Here, α = 1 corresponds to the Lasso penalty and α = 0 corresponds to the Ridge penalty. The term within the outer square brackets represents the elastic net penalty. In the context of this study, the predictor variables x_{ij} would correspond to the methylation level of the j-th CpG site, y_i would correspond to the age of the i-th individual, and the β_j’s are the coefficients that represent the contribution of each CpG site to the predicted age. The alpha is set to 0.5, and the 5-fold cross-validation was used to determine the optimal lambda. The model was trained using the training data, and the model performance was evaluated using the test data. The model performance was evaluated using the mean absolute error (MAE) and the root mean squared error (RMSE).

To obtain a single summary measure of biological age based on all Ageome clock predictors, we used the reversed-RMSE-weighted average of the Ageome clock predictions to calculate the MetroAge score:

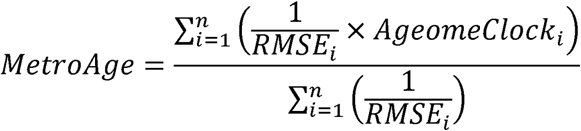

Each pathway’s age prediction is weighted by the inverse of its RMSE, giving more weight to the pathways that have lower errors (thus more accurate predictions). The sum of these weighted age predictions is then divided by the sum of the weights (the inverses of the RMSEs) to calculate the MetroAge score. The MetroAge score, therefore, is a measure of biological age that considers the accuracy of each pathway’s age prediction. Note that the RMSE here is calculated from cross-validation inside of the training data, therefore there is no data leakage.

### Pathway-specific Scaling Law Analysis

To investigate the evolutionary conservation of age-related epigenetic changes across species and functional pathways, we performed a scaling law analysis of pathway-specific methylation rates in mammals with varying maximum lifespans, extending the methodology described by Crofts et al., 2023 ^46^. We utilized DNA methylation data from 42 mammalian species, focusing on CpG sites that could be mapped across species.

For each pathway, we first filtered to include only those with at least 250 CpG sites to ensure robust analysis. We then applied the cumulative pairwise algorithm detailed in Crofts et al. (2023) to compare methylation slopes across species, using the mammal with the shortest observed lifespan (rat) as a reference. The methylation rate for the rat was set to 1, and rates for other species were calculated relative to this baseline. We computed scaling laws for each pathway using linear regression on a log-log scale, where the x-axis represented the maximum lifespan of different mammalian species and the y-axis displayed the relative methylation rate. The slope of this regression line represents the scaling law for each pathway.

To assess the probability of observing a scaling law under the null hypothesis that there are no pathway-specific effects, we created 100 random CpG pathways for each fixed size of pathway lengths (n=500, 2000, 1000, 500, and 250). We then computed the scaling law for each pathway using the exact same methodology (Extended Data Figure 1a). We fitted a normal distribution to the bootstrapped scaling laws for each fixed size (n) and observed that while the mean of the inferred scaling laws was constant, the spread or standard deviation of scaling law values (SD) correlated negatively with the size of the pathways (in the log-log scale, r2=-0.98, p=0.003, Extended Data Figure 1b). We use the predicted dynamics for mean and standard deviation, *m_n_*_sites_ and *S_n_*_sites_ as a function of the number of sites in a pathway, *n*_sites_. We then computed the two-tailed probability of observing a given scaling law x under the null hypothesis, or p-value, as

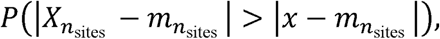

where, *X_n_*_sites_ ∼ *N*(*m_n_*_sites_, *s_n_*_sites_). We finally corrected the resulting p-value by the number of tested scaling laws to compute the FDR.

### Mortality analysis

Mortality analysis was performed in the Normative Aging Study (NAS) cohort (N = 1,488, 38.8% deceased), with DNA methylation data generated using the 450K array. The Biolearn framework was used to perform survival analysis using each Ageome clock model as the predictor ^28^. The hazard ratios (HR) and 95% confidence intervals (CI) were calculated for each Ageome clock. Cox proportional-hazards model ^47^ was used to test the association between each Ageome clock and the survival time. Chronological age is used as the covariate. The clock predictions were standardized before input into the model. The P-values were corrected for multiple testing using the Bonferroni correction.

### Disease association analysis

The MGB cohort consists of 4,246 subjects from the Mass General Brigham Biobank ^48^, with DNA methylation data generated using the EPICv1 array. Additionally, these subjects were linked to the Research Patient Data Repository to curate Electronic Medical Records (EMR) data. Consequently, 40 diseases were identified from the longitudinal diagnosis records using ICD codes (refer to the “Disease Codebook” Excel sheet, Extended table 1). Prevalent and incident cases were determined based on the chronological order of the first diagnosis record for a specific disease and the date of biosample collection. The time span was calculated accordingly, with negative values indicating diseases developed before biosample collection, used in the reverse test as occurrences of events, and positive values indicating the onset of new diseases after biosample collection, used in the forward test as events. Specifically, prevalent cases were excluded from the forward test. The Cox proportional-hazards model was applied to both forward and reverse tests, adjusting for age and sex in every model. To efficiently compute these numerous models, the R package RegParallel was utilized for parallel computation ^49^. All analyses were conducted using R 4.3.0.

## Acknowledgments

This work is supported by NIA and Hevolution grants. K.Y. was supported by NIH F99AG088431. E.LC. is supported by a EHA Bilateral grant number ID: BCG-202209-02649. We extend our gratitude to the Biomarker of Aging Consortium and members of the Gladyshev laboratory

