## Supplementary Figure 1 for "High-dimensional Ageome Representations of Biological Aging across Functional Modules"

Supplementary Data

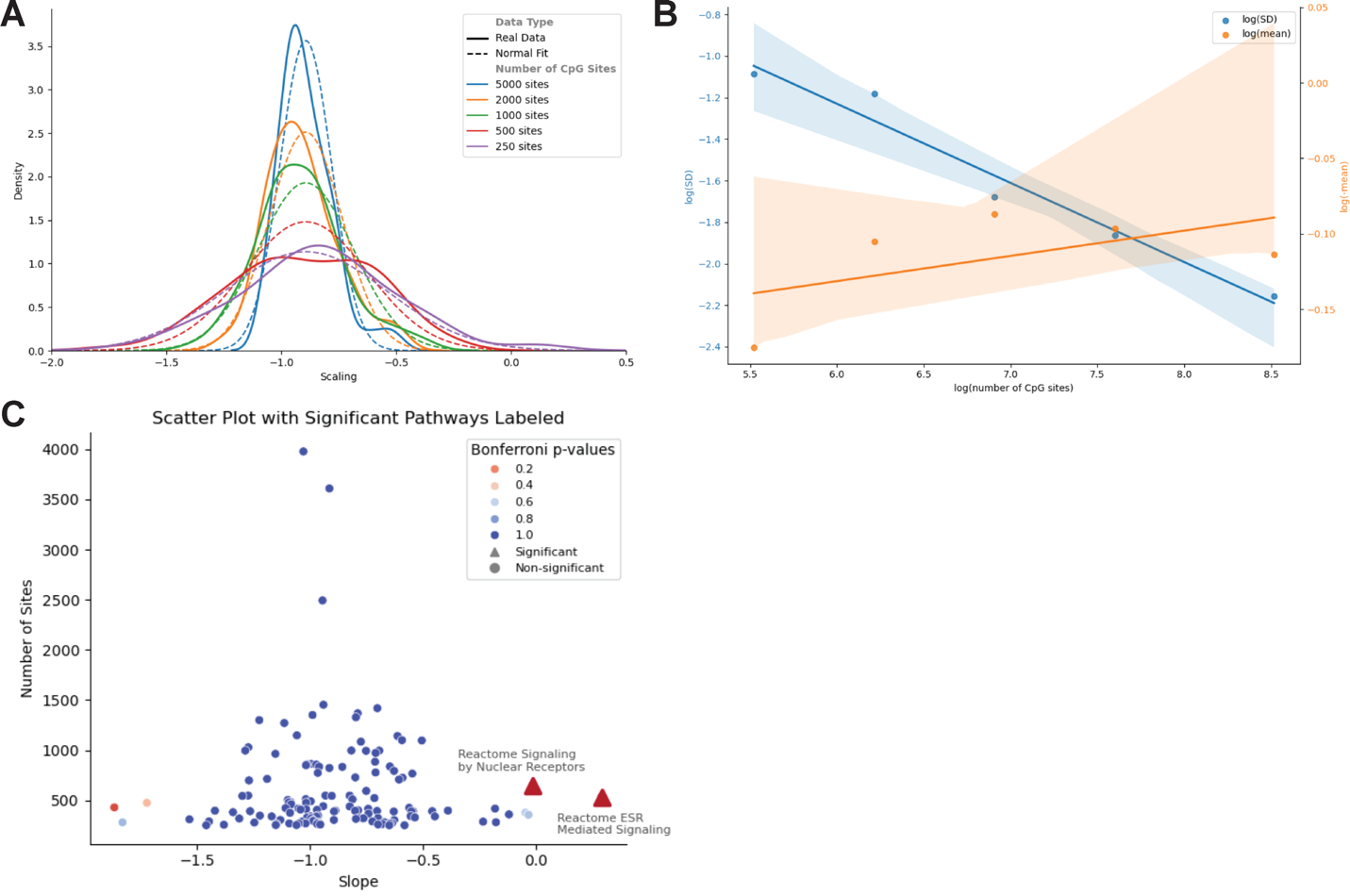

**Figure S1: Simulating the null hypothesis for methylation rate scaling laws**

(A) Simulation of the null hypothesis. Random subsets of CpGs with fixed sizes were created to simulate pathways. The scaling law for each random pathway was computed, and the resulting distribution of scaling laws was approximated using both Kernel Density Estimation (KDE) and a normal distribution.

(B) Description of mean and standard deviation (SD) of the normal approximation of scaling law distributions conditional on the number of CpG sites in the pathway.

(C) Probability of observing a pathway conditional on the number of CpG sites under the null hypothesis. The normal distribution description of scaling laws from simulated pathways was used to compute the probability of observing any pathway under the null hypothesis. Each point represents a pathway's number of CpG sites and inferred scaling law. Color scale indicates the Bonferroni-corrected p-values associated with the two-tailed probability of observing this scaling law conditional on the pathway's number of sites.

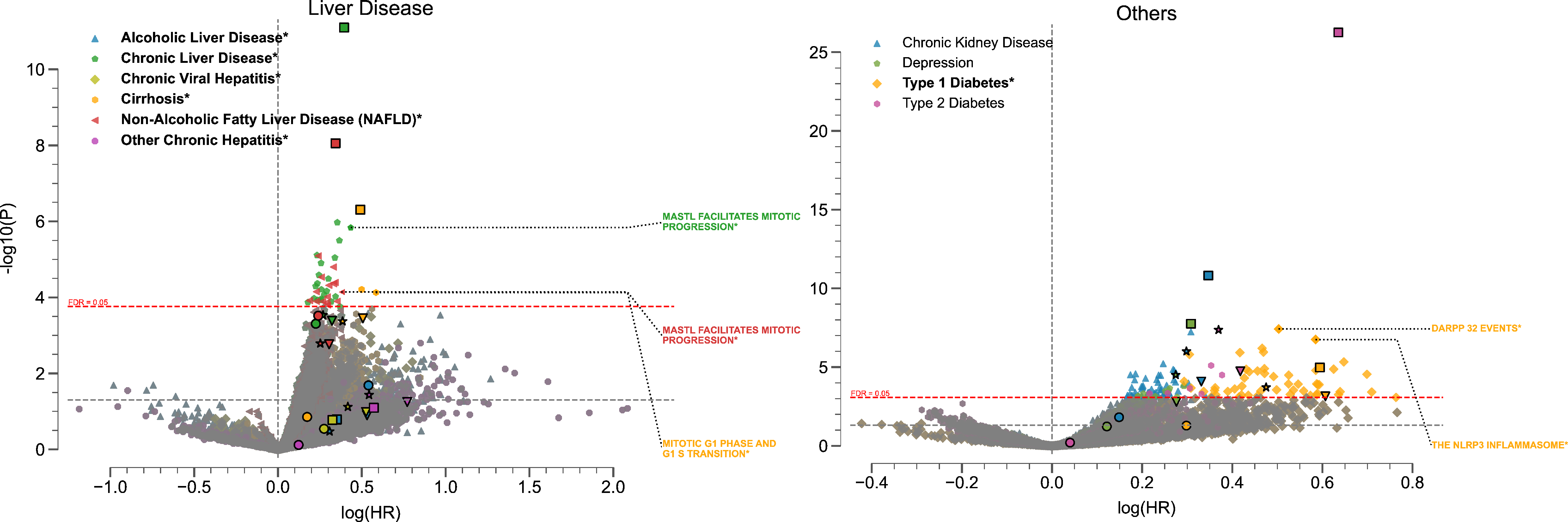

**Figure S2: Ageome predicts disease risk**

Volcano plots showing the relationship between log(HR) and -log10(P) for Ageome clock predictions of liver diseases (left) and other diseases (right). Top two pathways for each disease with P < 0.05 are labeled. Reference models are indicated by distinct markers with black borders: GrimAgeV2 (star), DunedinPACE (square), PhenoAge (triangle), and YingDamAge (circle). FDR threshold is set to 0.05, indicated by the dashed red line. The diseases that have ageome measurements that outperform (either by significance or hazard ratio) reference models are bolded and indicated by an asterisk (*), as well as the outperforming pathways. Only the top 2 Ageome pathways (ranked by -log10(P)*  log(HR)) with FDR < 0.05 are shown for each disease.

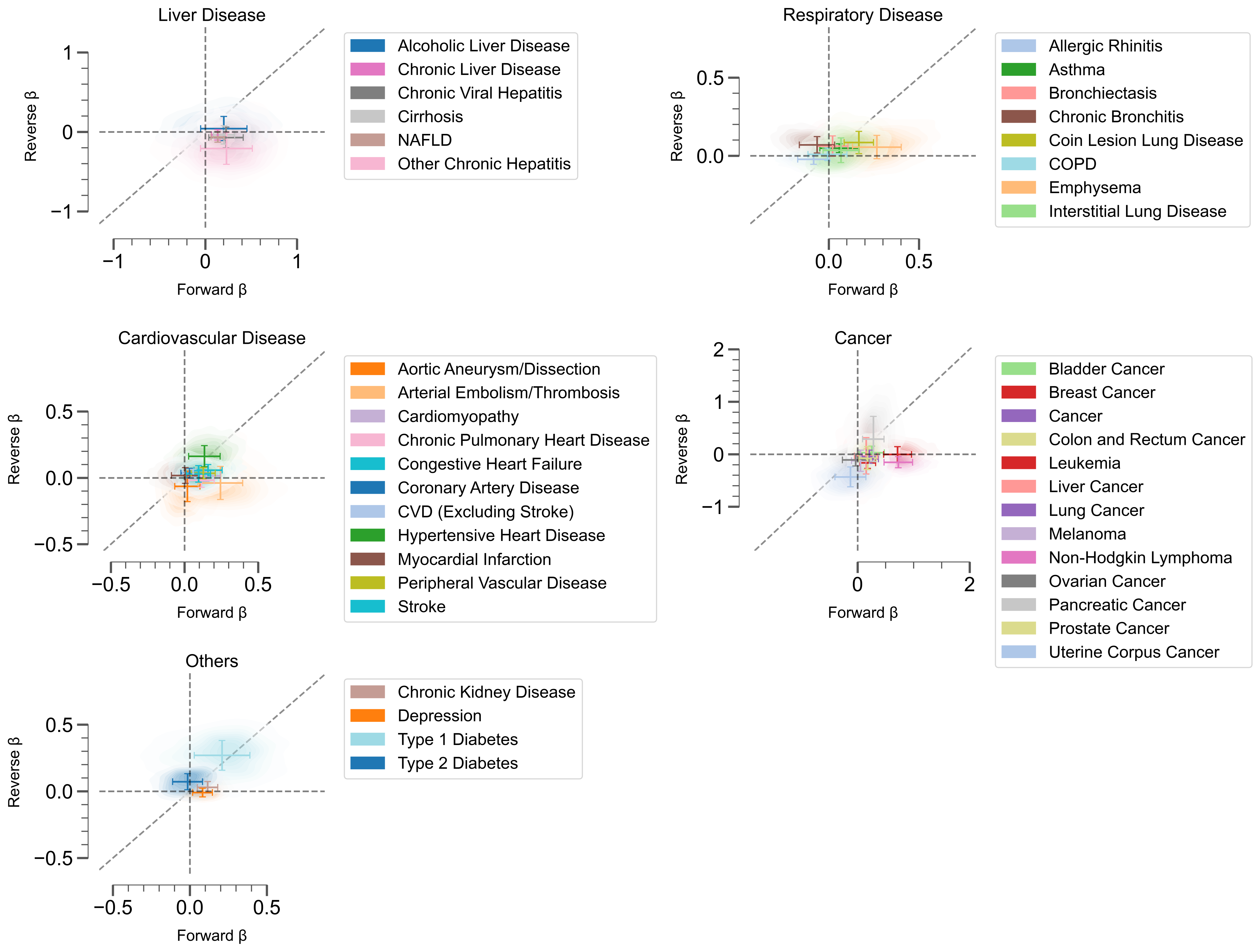

**Figure S3: Bidirectional analysis of aging-disease relationships in Ageome**

Scatter plot and 2D density plot shows forward (X-axis) and reverse (Y-axis) effects for different diseases. Error bar shows the standard deviation of the forward and reverse effects across Ageome clock measurements. The black dashed line shows the line where Δβ = 0.

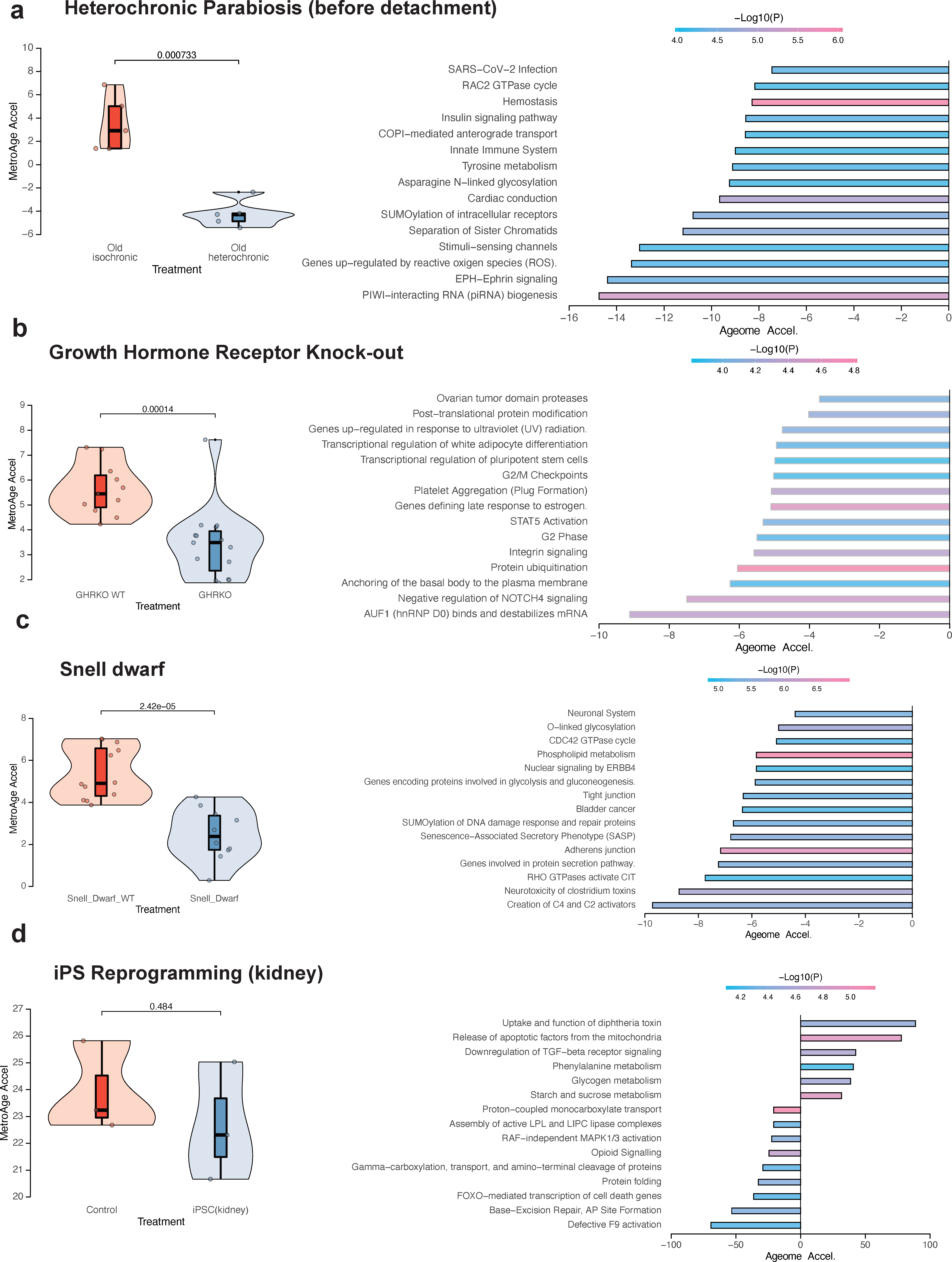

**Figure S4: Application of Ageome to various interventions**

Application of Ageome to heterochronic parabiosis model before detachment (A), growth hormone receptor knock-out (B), snell dwarf (C), and iPS reprogramming of kidney fibroblast (D). MetroAge indicates the overall effect of interventions (left). The bar plot highlights the top affected pathways based on each Ageome model (right). The color of the bars shows the -log10(P-value) of the Ageome for given interventions.
